# The Compositional Encoding of Hand-Eye Coordinated Movements for Single Neurons in the Posterior Parietal Cortex

**DOI:** 10.64898/2026.04.04.716384

**Authors:** Nikos A. Mynhier, Jorge Gamez, Kelsie Pejsa, Ausaf Bari, Richard M. Murray, Richard A. Andersen

## Abstract

Human posterior parietal cortex (PPC) is thought to play an important role in hand–eye coordination, yet the underlying encoding mechanisms remain uncertain. We recorded 412 single neurons across 11 sessions from motor cortex (MC; n=251) and PPC (n=161) in a single human participant performing a hand-eye (H-E) coordinated center-out task. While MC neurons showed little to no modulation by eye movements, 79% of PPC neurons had neural representations that were additively separable into independent hand- and eye-movement tuning curves. Due to this separability, neural representations could be separated and additively recomposed while maintaining structure similarity. Consequently, compositional decoders trained solely on single-effector movements could match the performance of decoders trained on coordinated H-E movements (hand: 66% vs 69%; eye: 34% vs 36%). These results show that, during simple center-out tasks, MC hand movement codes are unaffected by eye movements and that compositionality can be used to modularly decode H-E coordinated movements in PPC.

## INTRODUCTION

The human Posterior Parietal Cortex (PPC) is a core part of the dorsal ventral stream, a processing pathway that governs how vision is used for physical actions [1]. In particular, PPC encodes spatial processing for visually guided actions [2–4] and supports effective hand–eye (H-E) coordination during natural behaviors including reaching and grasping [5–7]. The encoding of hand-eye coordination in PPC has been studied extensively in nonhuman primate (NHP) models, with a wide variety of experimental paradigms [5–25]. However, throughout extensive study in NHPs, PPC has proved to be a heterogeneous region with many different areas and functional properties [26–30]. Additionally, many of these findings from monkey models have yet to be directly verified in humans. Instead lesion studies in human patients provide indirect confirmation that PPC plays a critical role in hand-eye coordination [31,32]. Damage to PPC can result in optic ataxia, a disorder characterized by impaired visually guided reaching despite intact primary visual and motor function. Patients with optic ataxia often misreach for targets, particularly in peripheral vision or contralesional space, consistent with a disruption in transforming visual information into appropriate motor commands [31]. PPC lesions can also produce spatial neglect syndromes, in which movements toward one side of space are systematically impaired [33]. Together, these deficits indicate that the link between vision and action are fundamentally compromised following PPC damage.

Hand and eye movements performed separately have previously been decoded from PPC in Brain-Machine Interface (BMI) studies [34,35]. However, the decoding of simultaneous hand and eye movements has yet to be explored systematically with BMIs. It is particularly important to be able to decode coordinated hand and eye movements because they commonly arise in BMI settings due to gaze shifts accompanying nearly all natural motor behaviors [36]. Although, the best way to decode simultaneous H-E movements is still unknown.

In recent years, BMI studies in the motor cortex (MC) have increasing observed the neural representations of multi-variable motor movements, such as multi-limb, multi-finger, and bimanual movements, to be compositional [37–42]. A neural representation is considered compositional when it can be separated into independent codes corresponding to each variable of a movement. Such structure has enabled the construction of complex BMIs which require multiple degrees of freedom such as writing or piloting cars and drones [43–45]. While motor movements appear to be compositional, it is unclear whether visual signals can also be composed with motor signals during H-E coordination. Moreover, while compositionality has been shown in PPC [46,47], most compositional BMIs rely on MC. Thus, here we test whether H-E representations in PPC exhibit the compositional structure and can support compositional decoding.

In this paper, we characterize single-neuron representations of hand–eye coordinated movements in the hand knob of MC and the superior parietal lobule (SPL) of SPL during a standard center-out task. In this context, we find that mixed selective neurons have additively separable codes for hand and eye movements. We also find that univariate (or single-effector) representations of isolated hand and eye movements generalize to multi-effector representations of coordinated hand-eye movements. Crucially, these properties allow us to create compositional decoders. We demonstrate that compositional decoders trained solely on single-effector data can predict coordinated hand–eye movements as well as decoders trained directly on multi-effector data. Together, these findings support that, in common BMI contexts, hand–eye representations in human SPL can be treated as compositional and support scalable, efficient decoding.

## RESULTS

### Single Neurons have composable and separable hand-eye representations

We performed a hand-eye coordination task with two kinds of trials, single-effector (SE) and multi-effector (ME) across 11 clinical sessions (Vid. S1). SE trials show neural responses to moving the hand and eye alone. ME trials show neural responses to moving the hand and the eye at the same time. Both kinds of trials use a 6-target center out structure. During an ME trial (Fig. 1A), both a hand reach and gaze shift are made simultaneously toward targets presented at angles θ_H_ and θ_E_. During the SE trials (Fig. 1D), only one movement (hand or eye) is made toward a single presented target with the other effector fixed (Vid. S2). During a run of the full task, SE and ME trials were pseudo-randomly interleaved, ensuring 3 repetitions of every possible target combination. Over the course of a session, 3 runs of the full task were completed (see Methods).

**Figure 1:**
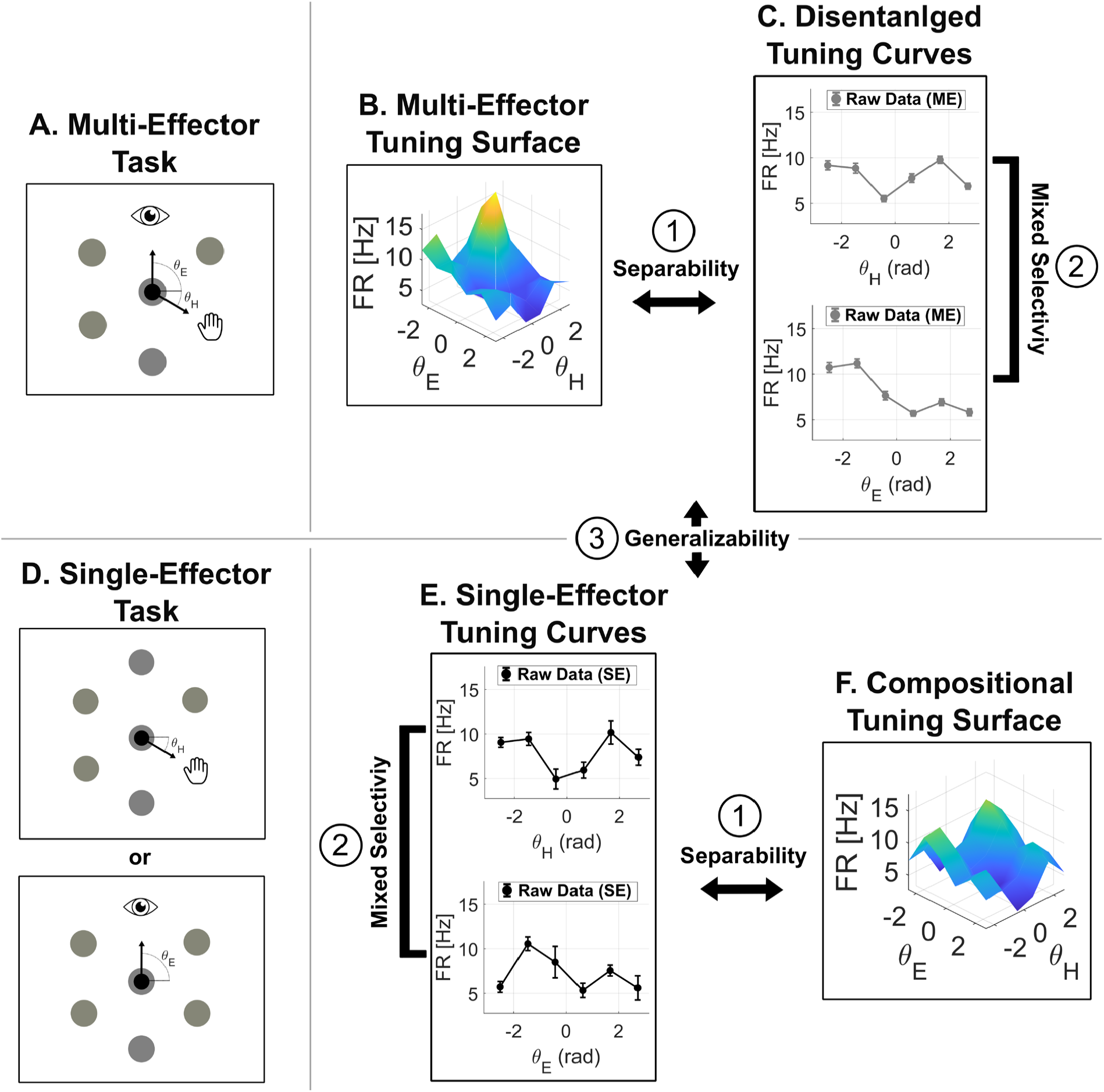
Single-session overview of task, tuning estimates, and decoding pipeline. (A) Schematic of the ME center-out task: two simultaneous targets (hand and eye) are presented at angular locations θ_H_ and θ_E_. Panels B,C,E,F are representations from an example neuron from the SPL of PPC. (B) Example SPL neuron’s ME tuning surface—firing rate (FR) as a function of hand and eye angles (θ_H_, θ_E_); surface color encodes FR [Hz] magnitude. (C) Tuning curves derived from the ME tuning surface (projections along θ_H_ and θ_E_). Points are means of the raw FR at each angle; error bars are standard errors. (D) Schematic of the SE tasks: the same hand and eye movements are performed on separate trials. (E) Tuning curves from SE trials (hand, eye), plotted as mean ± standard error of FR. (F) Additive SE surface formed by composing the two SE tuning curves (hand + eye + constant) on the (θ_H_,θ_E_) grid.

SE trials were a controlled characterization of how a neuron’s firing rate (FR) is modulated by the movement of only a single effector. The representation of an example neuron from SPL is presented as a discrete tuning curve (Fig. 1E). Tuning curves 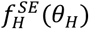 and 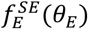 are presented as 6x1 vectors that describe the mean firing rate of the example neuron at each of the 6 discrete target angles (see Methods). ME trials characterized how the same example neuron’s FR was modulated by the simultaneous movement of the hand and eye. ME representations of this took the form of a 6×6 grid/tuning surface 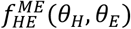 (Fig. 1B). This grid was created by spanning the combinations of the 6 hand and 6 eye movements (See Methods).

For this example neuron, ME surfaces 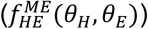 were able to be separated into one-dimensional tuning curves, 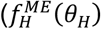 and 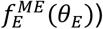 (Fig. 1C) (See Methods). Conversely, its SE tuning curves 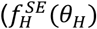 & 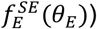 were able to be additively composed into a two-dimensional tuning surface 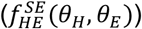 (Fig. 1F) (See Methods). Together, these results provide a conceptual framework for how joint hand-eye representations can be disentangled and equivalently how separate hand and eye representations can be additively composed.

### Single neuron representations are additively separable

A tuning surface is additively separable (*F*(θ_H_, θ_E_) = *f*(θ_H_) + *f*(θ_E_)) if and only if it has no cross-partial derivatives (∂^2^*F*(θ_H_, θ_E_)/∂θ_H_ ∂θ_E_ = 0) (see Methods). In other words, tuning surfaces separate into independent components when the encoded variables do not interact. Tuning surface cross-derivatives were calculated with centered wrap-around finite difference calculations (see Methods). Every neuron in MC and SPL was subject to a separability test using the cross-partial derivative of its tuning surface (Hz/rad^2^) (see Methods). The q-values for all neurons in both areas, arising from the separability test, create two distributions (Fig. 2A&2H). Because neurons in MC do not commonly encode eye movements, they should be trivially separable (*F*(θ_H_, θ_E_) = *f*(θ_H_) + 0). Consistent with that, all 251 (100%) neurons in MC were found to have no significant cross-derivative energy and thus were additively separable. In SPL, 127 (79%) neurons were found to have no significant cross derivative energy, while 34 (21%) neurons were found to have tuning surfaces with cross-derivative energy greater than a random surface. While a subset of neurons in SPL had significant variable interactions, it is possible to treat them as approximately separable if the strength of the interaction curvature is smaller than the curvature created from the main effectors.

**Figure 2:**
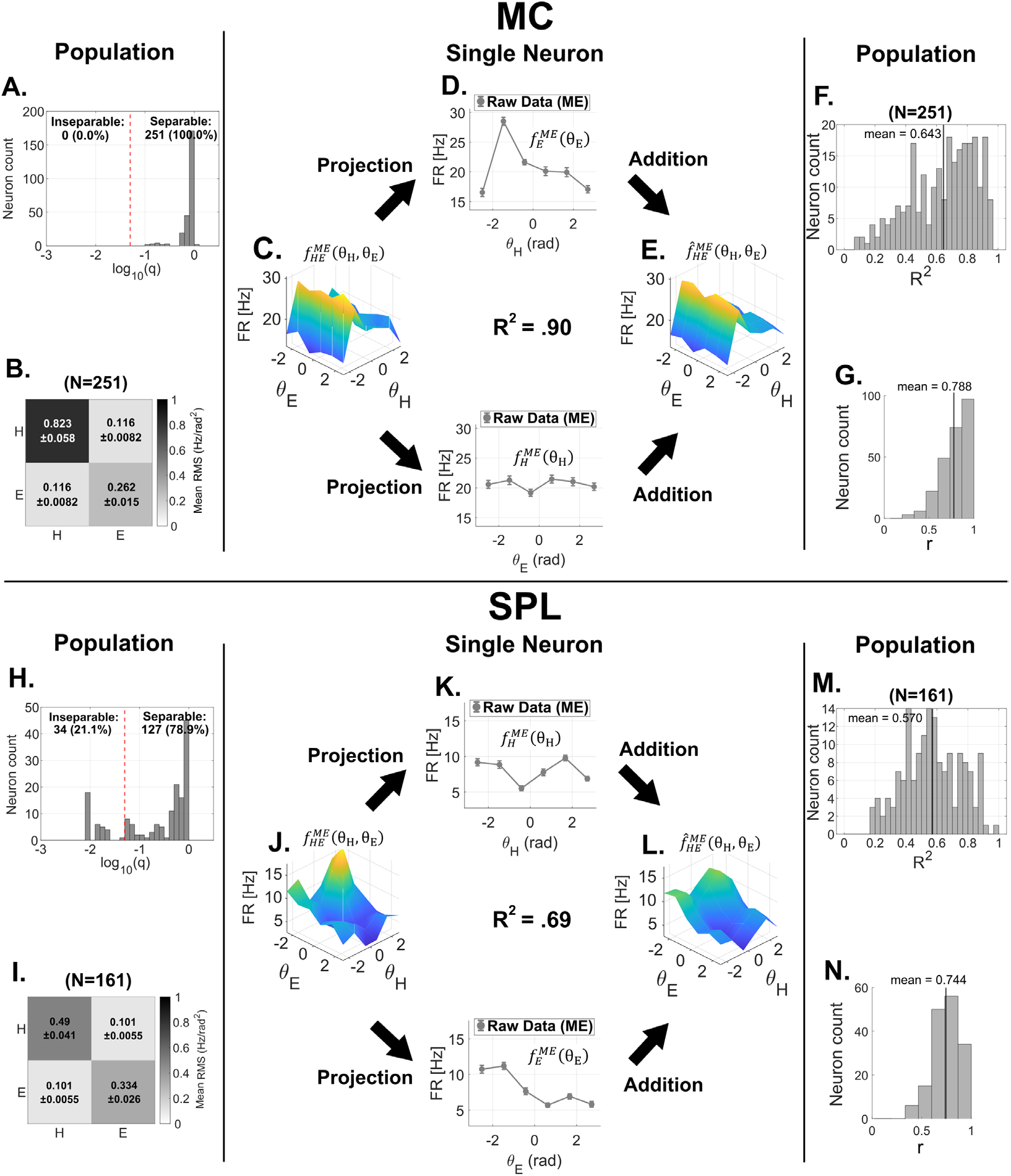
Separability of individual neurons during ME task. The figure tests the “separability” requirement of our compositional framework by comparing each neuron’s tuning surface during the multi-effector task to an additive reconstruction obtained from its projections and by quantifying cross-variable interaction. Panels A-G summarize the separability of neurons in the MC while H-N summarize neurons in the SPL. (A) Histogram of log(q) values (logarithm of FDR corrected p-values) arising from separability test across all neurons in MC. Significant q-values (left of the red significance threshold (*α* = . 05), imply the cross-partial derivative RMS of the raw surface is significantly greater than a random surface. (B) Heatmap showing the average value of the Hessians calculated for across every neuron’s tuning surface. The value of each element of the hessian summarizes the curvature of firing rate with respect to hand and eye angles. Off-diagonal entries characterize the interactions of the behavioral variables. Error bars show mean ± SEM across neurons. (C) Example neuron’s tuning surface 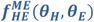 from the multi-effector task. Axes: hand direction θ_H_ (x), eye direction θ_E_ (y), firing rate (z; color also encodes FR). Directions are sampled on a 6×6 grid. (D) Projections of the tuning surface in panel C: hand tuning 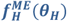 (top) and eye tuning 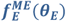 (bottom), obtained by averaging across the other effector. Points show means across trials; error bars are s.e.m. (E) Additive reconstruction 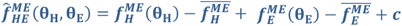, where 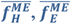 are the means of the tuning curves and c is a neuron-specific baseline estimated from the grand mean of the ME data. Axes and scale match panel C. (F) Distribution (histogram) of per-neuron reconstruction quality, reported as the coefficient of determination (R^2^) between the raw surface (in panel C) and the additive reconstruction (in panel E) . The total number of neurons contributing to the histogram is indicated on the panel (e.g., N=251). (G) Distribution (histogram) of per-neuron Pearson correlation coefficient between raw surface (panel C) and the additive reconstructed surface (panel E) with the same number of neurons. (H–N) are the same visuals as (A–G) but for the second cortical area, SPL.

Hessian matrices capture the local curvature of a function by quantifying its second-order partial derivatives with respect to a pair of variables. If a tuning surfaces’ Hessian is diagonal over its whole domain, it has no cross-partial derivatives and therefore is additively separable. In the context of neuronal representations, each element of a 2×2 Hessian reflects how sharply a neuron’s firing rate changes with respect to hand and eye movements, either independently (diagonal terms) or jointly (off-diagonal terms). (see Methods) Across all neurons in the MC, the average value of the Hessian’s symmetric off-diagonal terms was 0.12 ± 0.01 [Hz/rad^2^], while the diagonal terms were 0.82 ± 0.16 [Hz/rad^2^] and 0.26 ± 0.02 [Hz/rad^2^] for the second derivative of the hand and eye, respectively (Fig. 2B). Similarly for all neurons in the SPL, the average value of the off-diagonal terms was 0. 10 ± 0.01 [Hz/rad^2^] and the diagonal terms were 0.49 ± 0.04 [Hz/rad^2^] and 0.33 ± 0.02 [Hz/rad^2^] for the second derivative of the hand and eye, respectively (Fig. 2I). The interaction curvature strength is 20% of the hand curvature strength and 30% of the eye curvature strength, as measured by RMS deviation. So, while 34 neurons in SPL were found to significant variable interactions, on average the interaction curvature is much weaker than primary effector curvature. Due to the dominance of the separable population and the weakness of the average variable interactions we treat all SPL neurons as approximately separable.

Separable tuning surfaces can be expressed as a simple sum of independent tuning curves 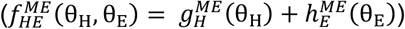. In the ME context, independent tuning curves for the hand 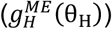 and eye 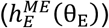 can be obtained by projecting the ME tuning surface onto one behavioral variable (averaging/marginalizing over the other variable) and centering. (see Methods) The ME tuning surface 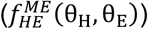 (Fig. 2C&J) and projected tuning curves 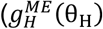 and 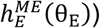 (Fig. 2D&K) are visualized for an example neuron from each brain area. For the representative neurons from MC and SPL, respectively, reconstructed additive surfaces 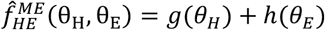 (Fig. 2E&L) well approximated the pre-projection empirical ME tuning surface (*R*^2^ = .9 and *R*^2^ = .69).

To quantify reconstruction quality across all neurons, we compared 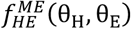 and 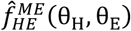 over the common 6×6 grid using a weighted coefficient of determination (see Methods). The mean of the per-neuron R^2^ distribution was .64 in MC and .57 in SPL, therefore the average additively reconstructed surfaces were moderately linearly related to their raw empirical counterparts (Fig. 2F&M). Across the distribution of all neurons the average Pearson correlation coefficients (*ρ*) between the raw and reconstructed tuning surfaces was .79 in MC and .74 in SPL, indicating a consistently positive and strong linear association between the surfaces of all neurons (Fig. 2G&N).

While the data suggest that neural representations are separable on the single neuron level it is common to investigate if separability extends to population codes. Conveniently, it is possible to analytically prove that if all neurons are additively separable then any linear projection of the population activity will also be separable. (see Methods) We tested this implication for each session and identified a demixed additive basis where the projected population activity was additively separable, as evidenced by a parallelogram structure in the population latent space (Supplemental Figure 1). The variety of measures reported here provide convergent empirical evidence that neurons in both brain regions can be reasonably treated as additively separable.

### Neurons in SPL are mixed selective

After establishing separability of single neuron representations, it is justified to treat projections of tuning surfaces as single variable tuning curves. Projected ME tuning curves and empirical SE tuning curves were used to calculate selectivity of all recorded neurons. Commonly, selectivity is determined by an ANOVA. However, we found the distribution of data for each trial condition to be heteroscedastic with non-Gaussian firing rate residuals thereby violating the assumptions of ANOVAs (Supplemental Fig. 2). This motivated the use of a nonparametric approach to assess tuning significance. The nonparametric approach we used was a permutation test of tuning-curve amplitude (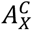 Fig. 3A&F), with significance determined from Benjamini–Hochberg–adjusted q-values 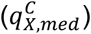 computed across bootstrap resamples (see Methods). Neurons were assigned to four classes by significance testing: “Hand” or “Eye” if only one effector’s tuning curve amplitude was significantly above baseline; “Both” if both were; and “None” if none were. This analysis revealed a marked difference in effector selectivity between the two cortical areas. In MC (Fig. 3B), the population was dominated by hand-selective neurons (≈85%), with very few eye-selective (<1%) or mixed-selective (∼1%) units, and a minority of non-selective neurons (≈14%). In contrast, SPL (Fig. 3G) exhibited a large population of mixed-selective neurons (≈50%), alongside hand-selective (≈39%), eye-selective (≈4%), and non-selective (≈7%) neurons.

**Figure 3:**
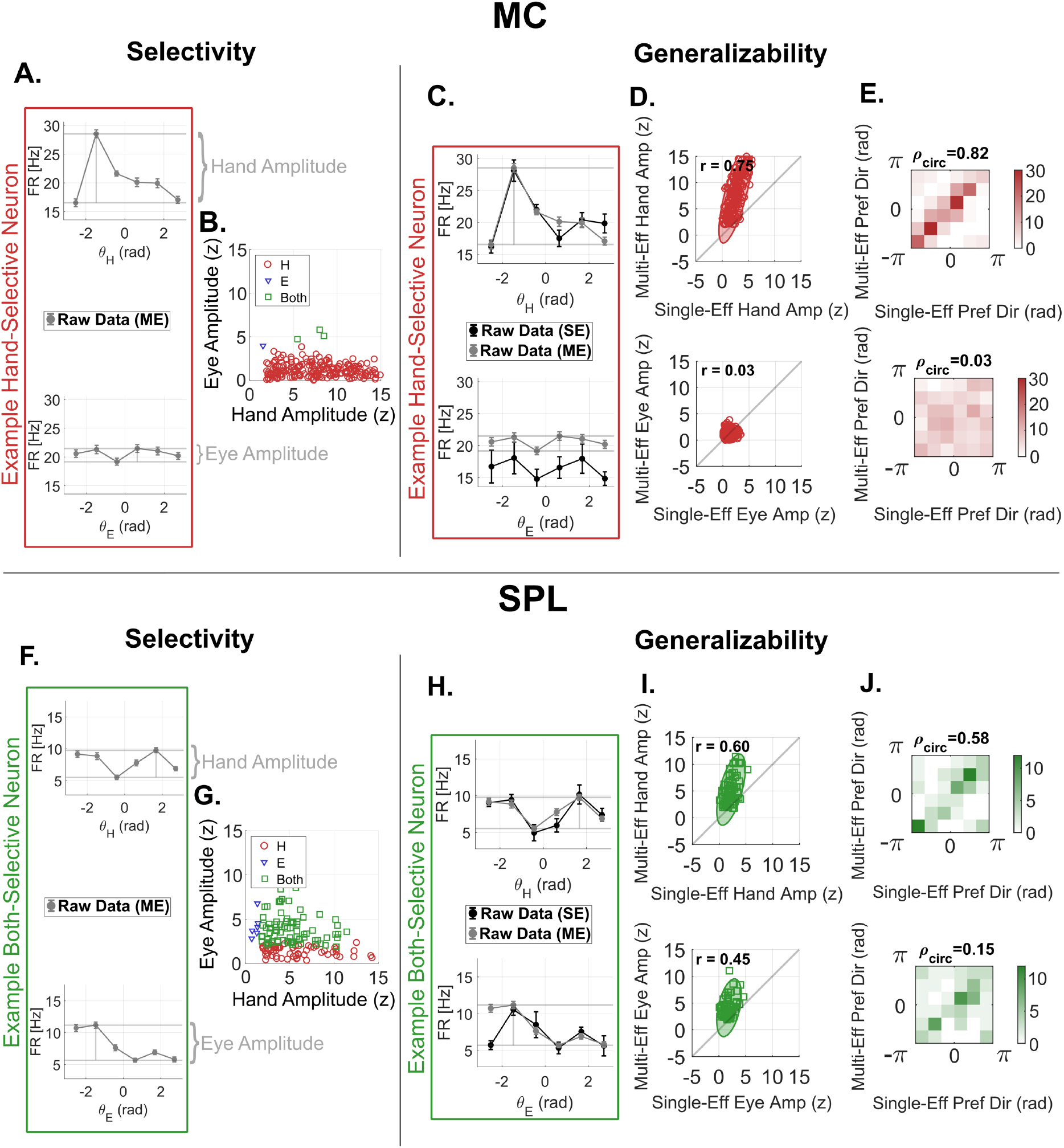
Selectivity and generalizability. This figure demonstrates the neuronal selectivity and generalizability properties for neurons with compositional coding. Selectivity is determined by testing tuning-curve amplitudes against a permutation-based null; neurons significant for hand, eye, or both are labeled H, E, or Both. Correlation in tuning amplitude and preferred direction (angular bin with maximum mean response) are used to demonstrate generalizability. (A–E) Motor Cortex (MC) data: (A) Example hand and eye tuning curves; a light gray segment mark tuning curve peak and trough with a vertical line centered at the tuning-curve preferred direction. (B) Population scatter of z-scored amplitudes (hand on x-axis, eye on y-axis) for all neurons; points are colored and shaped by significance outcome (red circle, H; blue triangle, E; green square, Both). (C) overlaid ME (gray) and SE (black) hand tuning curves for an example hand-selective neuron; (D) scatter of z-scored amplitudes in SE (x-axis) versus ME (y-axis) for hand-selective neurons, with Pearson r reported; (E) confusion matrix comparing SE and ME preferred directions for the same neurons. (F–J) share a parallel panel structure to A-E but for the SPL. (F) amplitude definition on example tuning curves; (G) population amplitude scatter with the same color code; (H) overlaid ME (gray) and SE (black) tuning curves for an example neuron categorized as Both (green outline); (I) SE–ME amplitude scatter for Both-selective neurons, Pearson r reported; (J) confusion matrix comparing SE and ME preferred directions for the same Both-selective neurons. All amplitudes are computed relative to baseline and z-scored using the permutation-derived null.

### Neuronal representations are generalizable across experimental contexts

To assess generalizability of the representations across ME and SE contexts, projected ME tuning curves 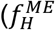 & 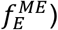 were compared to empirical SE tuning curves 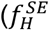 & 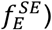in both MC (Fig. 3C) and SPL (Fig.3H). Two metrics were used for comparison, tuning curve peak-to-peak amplitude 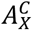 (Fig. 3D&I) and preferred direction 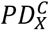 (Fig. 3E&J). (See Methods)

In the MC, hand-selective neurons (making up ≈85% of MC neurons) demonstrated hand tuning curves with highly correlated amplitudes and PDs between the ME and SE contexts. (r=.75 and r=.82 respectively) (Fig. 3D&E) In the SPL, mixed selective neurons (making up ≈50% of SPL neurons) demonstrated preservation of both hand and eye representations across contexts. Hand tuning curves showed high correlation of amplitudes and PDs across contexts (r=.60 and r=.48 respectively). Eye tuning curves showed moderate correlation of amplitudes (r=.45) but weaker correlation of PD (r= .15) across contexts. While present in SPL, eye representations were generally weaker and noisier than hand representations.

Projected ME tuning curves have a baseline FR equal to the grand mean of the ME tuning surface, which is affected by both hand and eye movements. SE tuning curves, which involve movement of only one variable, have baseline FRs that differ from those of coordinated movements. This discrepancy is most apparent for eye tuning curves in MC (Fig. 3C). Because eye movements are not encoded in MC, the SE eye tuning curve baseline is low, whereas the projected ME eye tuning curve baseline is elevated due to the influence of concurrent hand movements. When composing SE tuning curves into surfaces we accounted for this by calculating the predicted baseline FR for the tuning surface as a linear combination of the SE baselines (see Methods).

Despite being well correlated, ME tuning curves showed systematically higher z-scored peak-to-peak amplitudes than their SE counterparts (Fig. 3D&I). This is caused by greater variation in the SE tuning curve amplitudes, in part due to a fewer number of SE trials. Across 251 neurons in the MC, hand tuning curve amplitudes had an average coefficient of variation (CV) of .15 for ME data but .24 for SE data across the. Across 161 neurons in the SPL, hand tuning curve amplitudes had an average CV of .20 for ME data and .26 for SE data. This systematic difference in CV resulted in a greater-than-one ratio between ME and SE z-scored amplitudes across neurons in both brain areas. (Fig. 3D&I).

Together these results show that SPL contains many mixed-selective neurons whose hand and eye tunings are both generalizable from the SE to ME context. The SPL population is therefore separable, mixed selective and generalizable. Subsequently, these properties can be leveraged to build compositional, SE– derived generative models capable of decoding ME movements.

### Composed single-effector models decode multi-effector behaviors

After establishing that a subset of PPC neurons were (i) additively separable, (ii) mixed-selective, and (iii) generalizable, it was possible to create compositional surfaces from SE data that are comparable to the raw ME surfaces. If the two surfaces are effectively the same, from the perspective of likelihood evaluation, their decoding performance should also coincide.

Using the ME data (Fig. 4A&B), we estimated each neuron’s tuning surface 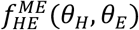 from 216 training trials (36 hand–eye combinations × 6 repeats). For the SE data (Fig. 4C&D), we estimated the compositional tuning surface 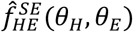 by summing two SE tuning curves per neuron 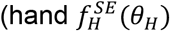 and eye 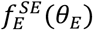 estimated from 108 SE training trials (6 directions × 9 repeats x 2 effectors). Three similarity metrics (i) Pearson correlation, (ii) grand mean FR, and (iii) peak FR, all showed that the SE-based compositional surface is approximately equal to the raw ME tuning surface. (Fig. 4E-G)

**Figure 4.**
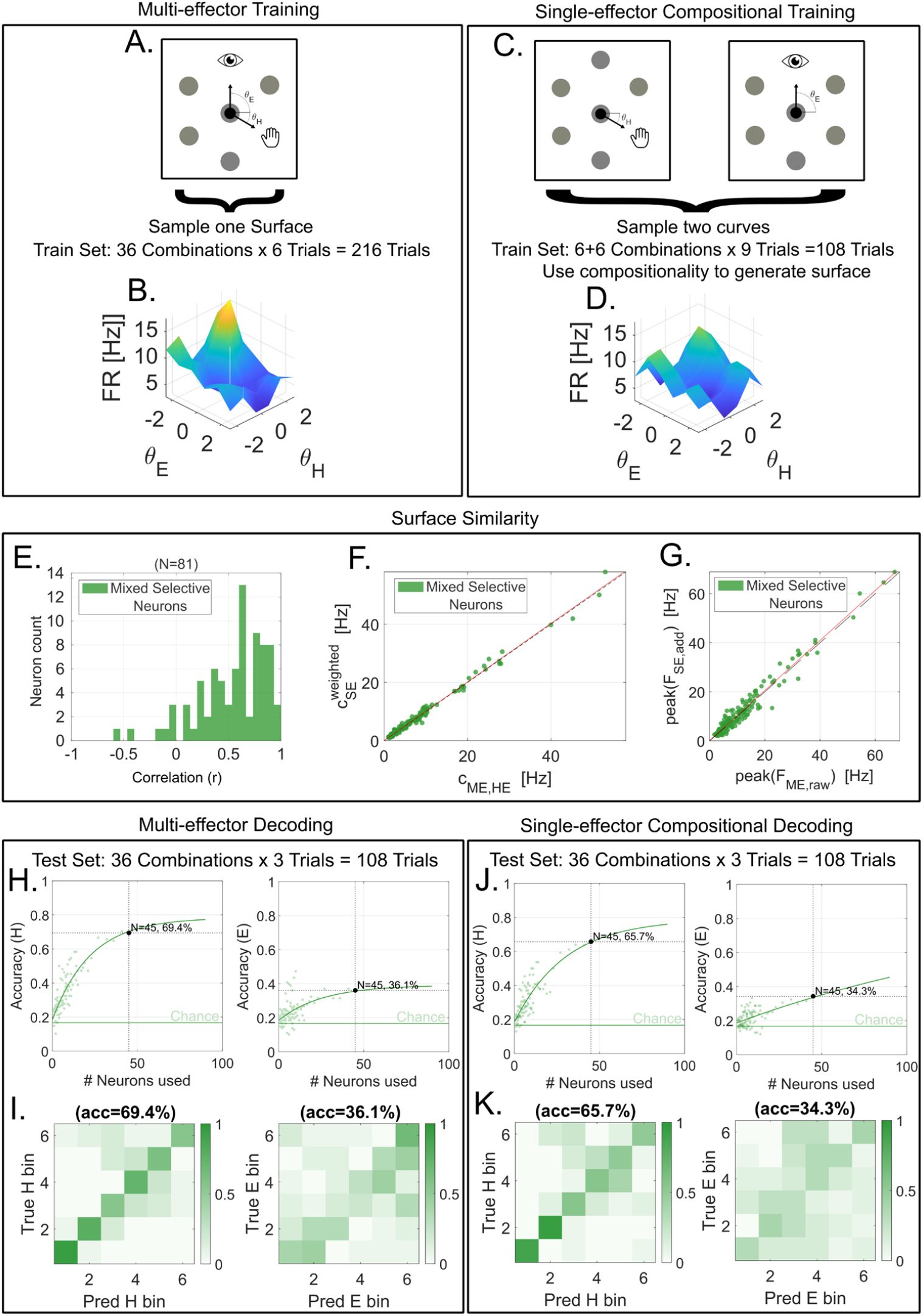
Compositional SE models decode as well as multi-effector models in SPL. All panels use posterior parietal cortex (SPL) data. (A) Schematic of the ME center-out task in which hand and eye targets at angles θ_H_ and θ_E_ are presented simultaneously. Training uses 216 trials per session (36 combinations × 6 repeats). (B) Example tuning surface f(θ_H_,θ_E_) derived from ME raw data (surface height and color encode z-scored firing rate). (C) Schematic of the SE (SE) tasks in which hand and eye movements are executed on separate trials in the same six directions; training uses 108 trials per session (hand: 6×9; eye: 6×9). (D) Additive compositional surface constructed from SE curves. (E) Histogram of Pearson correlation coefficients between ME and SE tuning surfaces for 81 mixed selective neurons in SPL. (F) Scatter plot of baseline FR for the raw ME surface versus additive SE surfaces of 81 both selective neurons. (G) Scatter plot of peak FR for the raw ME surface versus additive SE surface for same 81 neurons. (H) Inverse dropping curve: Decoding accuracy as a function of the number of neurons for hand (left) and eye (right); gray points are per-session accuracies and black curves are saturating fits extrapolated beyond the observed range. (I) Decoding confusion matrices for hand (left) and eye (right) from session with most neurons; rows are true bins, columns are predicted bins, values are row-normalized, and mean diagonal accuracy is shown above each matrix. (J) Inverse dropping curve but for the accuracy of prediction with the compositional model (format as in G; same ME test set). (K) Decoding confusion matrices for the compositional decoder (format as in H). Panels H-K enable a direct, like-for-like comparison between ME-trained and SE-trained decoding on a multi-effector (ME) test set of 108 trials per session (36 hand–eye combinations × 3 repeats).

For all surfaces we used circular kernel regression to obtain a smooth, continuous estimate of the mean firing rate at any behavioral state. (see Methods) Using this kernel we defined a likelihood *p*(*r*_*ref*_∣)*θ* for each surface given an observed firing rate (*r*_*ref*_). Assuming conditional independence of neurons we created a population likelihood by summing the log-likelihoods across neurons. We used the agreement of the independent neuron likelihoods in the population, as well as assuming a uniform prior over *Θ*, to perform Maximum a Posteriori (MAP) estimation of effector state. (see Methods) This enabled the creation of two MAP based decoders, one for the SE-based compositional surfaces and another for the raw ME surfaces. MAP decoding accuracies are presented as the fractions of test trials with correctly inferred hand (or eye) state (between 6 discrete bins accuracy from pure chance is 16.67%). MAP decoders accuracies were evaluated on an identical held out ME test set of 108 trials per session (36 combinations × 3 repeats).

Using an empirical maximum of 45 mixed-selective neurons, ME decoding accuracies for hand and eye movements were 69% and 36%, respectively, while SE decoding accuracies were 66% and 34%. Inverted dropping curves (Fig. 4H&J) further demonstrated near-identical accuracy scaling for ME- and SE-trained models, with saturating fits that extrapolated to similar ceilings (see Methods). However, with an average of approximately 15 neurons recorded during a single session the average decoding accuracy of hand and eye movements in ME trials were 38% and 25%, respectively, compared to 34% and 22% from the SE trials. (Fig. 4I&K)

Despite the ME model being trained with twice as many trials as the SE-based compositional model (216 trials vs. 108 trials), the compositional model achieved roughly the same decoding accuracy when evaluated on the same test set. The SE-based decoding succeeded precisely because the SE-composed surface 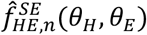 was an accurate surrogate for the ME surface 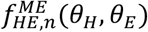. Therefore, the decoding results and representational similarity metrics are decisive evidence of the compositional coding of single neurons in PPC during the execution of a H-E coordinated center-out task.

## DISCUSSION

The central result of this paper is that, in the SPL of PPC, hand and eye movement codes are independent components of the neural representations arising from H-E coordinated movements during conventional center-out tasks. This is significant because it implies that in SPL eye movement activity can be disentangled or marginalized over during the decoding of certain coordinated behaviors. We also demonstrate that separable codes can be used compositionally to decode coordinated movements from models trained on isolated movements. In addition to the scientific results, this work also develops a more general methodology for identifying compositionality with just three necessary and sufficient representational properties.

The first property, additive separability, ensures that multi-effector (ME) tuning surfaces are well approximated by the sum of their one-dimensional projections (Fig. 2). To identify additive separability in neural representations, we use cross-derivative energy as diagnostic tool (Fig. 2A-B,H-I). This tool is particularly useful because zero cross derivative energy alone is a necessary and sufficient condition for additive separability (see Methods). Beyond hand–eye coordination, the cross-derivative framework also generalizes naturally: in *n* variables, additive separability corresponds to a diagonal Hessian over the full domain and the vanishing of all mixed finite differences. We also posit that this representational feature could be embedded into learning objectives by penalizing cross-derivative energy, thereby biasing models toward additive, compositional solutions.

The second property, mixed selectivity [48], establishes that the separable projections of tuning surfaces are significantly correlated to a behavioral variable. Here the contrast between SPL and MC was stark. MC neurons were overwhelmingly hand-tuned, with negligible encoding of eye movements (Fig. 3A,B). In contrast, approximately half of neurons in SPL were significantly tuned to both hand and eye (Fig. 3F,G). These findings are consistent with known differences in functional specialization between MC and SPL. The MC has many subregions, each of which encode the motor commands of specific body parts with few confounding signals. Meanwhile, the SPL encodes movement commands across many parts of the body, including the hand and eye, as well as potentially movement in a common distributed space [26,49].

The third and final property, generalizability, guarantees that independent neural codes are consistent across single- and multi-effector behavioral contexts. In SPL, the amplitude and preferred direction of hand movement representations were shown to be moderately to strongly correlated between SE and ME contexts. Meanwhile, for eye movement representations, amplitude and preferred direction were only moderately correlated between contexts, in part driven by weaker overall encoding strength (Fig. 3I,J). These results indicate that SPL provides stable, reusable representations of both hand and eye, with evident asymmetry in strength.

When these conditions hold simultaneously, models assembled solely from SE-derived modular tuning curves can be used to accurately decode ME behaviors. A compositional decoder trained solely on SE data achieved the same accuracy on held-out ME trials as a decoder trained directly on ME tuning surfaces, even though the latter had access to twice as many training samples (Fig. 4I-K). This equivalence demonstrates the advantages afforded by compositionality. The ability to build accurate ME decoders from reusable SE modules suggests that future BMIs could exploit compositional codes to reduce training demands and improve versatility of BMIs [40–42]. Moreover, this modular encoding strategy also facilitates the subtraction of unwanted separable neural codes thereby improving decoding performance in the presence of confounding behavioral variables. If similar compositional structure extends to additional effectors encoded in PPC, this region may represent a promising target for future BMI implantation.

Another contribution of this work is the demonstration of simple, nonparametric likelihood-based decoders. When making scientific deductions, decoding performance is commonly used as a proxy for representational structure in BMI studies [34,46]. However, likelihood-based decoders directly link representation geometry to decoding performance. This link shifts the construction of compositional decoders to the representational level, where additive neural components can be combined intuitively and then mapped to decoding.

While the findings presented here largely pertain to single neuron representations, we also proved that single neuron additive separability guarantees additive separability of population codes under linear projections. (See Methods and Supplemental Figure 1) Mixed selectivity and generalizability trivially extend to population codes as well. This is a particularly important revelation as it unifies single neuron- and population-level conclusions. However, it also emphasizes that at both the single neuron and population level, neurons in the SPL can be treated as additively compositional.

Taken together, these results demonstrate that, during certain H-E coordinated behaviors, the SPL of PPC employs low-interaction nearly independent representations of multiple effectors. More broadly, this work introduces a useful quantitative method for probing when neural systems operate in modular regimes. The joint presence of additive separability, mixed selectivity, and cross-context generalizability establishes a concrete, testable notion of compositionality. This demonstration of compositionality adds to growing evidence of how neural decoding can benefit from identifying intuitive components of neural representations. Compositionality provides a strong intuitive link between additively separable neural representations and modular decoders. From a translational perspective, this implies that accurate multi-effector BMIs can be constructed from modular codes learned in simpler settings.

### Limitations of the study

Several limitations of this study should be acknowledged. First, all data were obtained from a single participant, making this work a case study. While the findings provide a proof of principle that SPL neurons exhibit compositional coding of hand–eye movements, broader generalization will require replication in additional participants and in neighboring brain regions to establish whether the observed properties are consistent across individuals and cortical areas.

Second, the experimental design imposes constraints on how hand and eye representations can be dissociated. Because participants must use their eyes to perceive target cues, it is difficult to elicit hand movements independently of eye movements. Because of this, small eye movements are occasionally made over the course trials that are out of alignment with this goal/eye target. This coupling creates a potential confound, as eye activity is inextricably linked to the command structure of hand movements in this task. We did not observe any consistent behavioral confounds due to this task structure but it likely contributed to the noise floor of the experimental results. One possible solution is to study compositionality in naturalistic behaviors without explicit trial commands, where neural and behavioral activity can be analyzed post hoc.

Beyond these explicit constraints, several subtler issues warrant mention. The task design discretized both hand and eye movements into six directions, and all analyses were performed on the resulting 6 × 6 tuning grids. This coarse discretization may obscure finer-grained nonlinearities or interactions that only emerge in continuous movement spaces. Likewise, while the cross-derivative energy test is mathematically rigorous, finite differences on a sparse grid may lack sensitivity to local deviations from additivity. Thus, our conclusion of negligible interaction energy should be interpreted in light of the resolution limits imposed by the task and sampling.

A natural extension of this work is to characterize how the structure of neural representations depends on behavioral regime. Although representations appear largely separable in simple center-out tasks, more dynamic or naturalistic behaviors may induce stronger variable interactions. Determining which conditions favor separable versus interaction-dominated structure may sharpen our understanding of PPC function. It may also inform the design of future behavioral tasks which hope to specifically leverage separable codes.

A further limitation concerns the asymmetry of representations in SPL. While hand signals were robust, eye signals were consistently weaker and noisier. This asymmetry raises the possibility that neurons in the SPL only appear additive compositionality due to the weakness of the eye representation. If the eye is weakly represented, then it is possible that variable interactions with the eye are weak as well. Extending the analysis to effectors with more balanced encoding strength, such as bimanual limb movements, may clarify whether compositionality is a universal property of SPL or a result of asymmetric encoding.

Finally, the decoding analyses, while informative, rested on assumptions of conditional independence across neurons and stationarity of tuning over the course of recording sessions. Both assumptions are common in BMI modeling but are known simplifications of neural population dynamics. Dependencies across neurons or nonstationarities in tuning could alter the apparent degree of compositionality or its utility for decoding. Future work using larger-scale simultaneous recordings and models that explicitly account for neural correlations may provide a more complete picture.

Together, these limitations highlight the need for additional work to validate compositionality as a general coding principle. While the present study establishes a strong initial case in SPL, broader evidence across participants, brain areas, behavioral paradigms, and analytical frameworks will be essential to confirm its scope and robustness.

## RESOURCE AVAILABILITY

### Lead contact

Requests for further information and resources should be directed to and will be fulfilled by the lead contact, Nikos Mynhier.

## Materials availability

- This study did not generate new unique reagents.

## Data and code availability

- All data produced in the present study will be publicly available as of the date of publication.
- All original code produced in the present study will be publicly available as of the date of publication.
- Any additional information required to reanalyze the data reported in the paper is available from the lead contact upon request.

## ACKNOWLEDGMENTS

We thank participant RD for engaging in our clinical trial. We thank Viktor Scherbatyuk for IT and technical support. This work was funded by the National Eye Institute Award UG1EY032039A, the T&C Chen Brain-Machine Interface Center, and the James G. Boswell Foundation. We also would like to give special acknowledgement to Kellan P. Moorse for strongly contributing to the conceptualization of the project and to Sunho Lee and for consulting on mathematical formalism.

## AUTHOR CONTRIBUTIONS

Conceptualization, N.A.M., R.A.A., J.G., and R.M.M.; methodology, N.A.M., and J.G.; Investigation, N.A.M.; writing—original draft, N.A.M.; writing—review & editing, R.A.A., R.M.M., J.G.; funding acquisition, R.A.A., K.P., J.G., and N.A.M.; resources, R.A.A., K.P., & R.M.M.; Clinical Trial Management, K.P., R.A.A., J.G.; supervision, R.A.A., R.M.M., J.G.; Surgical and Medical Intervention, A.B.

## DECLARATION OF GENERATIVE AI AND AI-ASSISTED TECHNOLOGIES

During the preparation of this work, the author(s) used ChatGPT to proofread including the revision of grammar, syntax, and structure of written text. ChatGPT was also used to verify and condense code used for data analysis. After using this tool or service, the author(s) reviewed and edited the content as needed and take(s) full responsibility for the content of the publication.

## ADDITIONAL RESOURCES

This research is part of a human brain-machine interface clinical trial (NCT01958086): https://clinicaltrials.gov/study/NCT01958086?term=NCT01958086&rank=1

## STAR★METHODS

### EXPERIMENTAL MODEL AND STUDY PARTICIPANT DETAILS

A 40-year-old male participant (RD) was implanted with four 64-channel NeuroPort Utah electrode arrays. Four implant locations were determined using anatomical priors and preoperative fMRI (Supplemental Figure 3). The first array was implanted in the Superior Parietal Lobule (SPL) in the PPC. The second array was implanted in the Supramarginal Gyrus (SMG) of the PPC. SMG was found to not respond to hand and eye movements and was thus not used in this study. The third and fourth arrays (MCM and MCL) were implanted in medial and lateral regions of the motor cortex hand knob, respectively. Due to functional similarity, these arrays were pooled and analyzed as a single motor cortex (MC) population.

## METHOD DETAILS

### Experiment/Multi-effector Task

We developed a multi-effector center-out task (Fig. 1A&D; Vid. S1&S2) to induce hand-eye coordination. Participants were instructed to perform simultaneous reaches (hand movements) and gaze shifts (eye movements) toward two distinct targets displayed on a screen. Both targets were selected from a set of six possible radial locations. SE trials (e.g., only a reach or only a saccade) and rest trials (e.g. no movement) were also included as controls. The length of each trial was uniformly randomly sampled such that the rhythm of the task would not confound behavioral commands and preventing the participant from any anticipatory or preparatory movements. This resulted in a 1-1.5 second inter-trial interval and 2.5-3.5 second Go-phase where the movements were made. The movement angle, from the center of the screen, is the primary behavioral metric used for both hand and eye movements. To verify the participant’s correct execution of the task, eye movements were recorded with a 90 Hz binocular Tobii eye tracking system, and the hand movements were recorded using an external video camera. During one block of this task every possible combination of targets occurred 3 times. On each of the 11 recording sessions, the blocks were performed two or three times.

### Neural Signal Recording and Processing

Neural signals were recorded using the NeuroPort Neural Signal Processors (Blackrock Microsystems Inc.), then amplified, filtered with a bandpass range of 0.25 to 5 kHz, and digitized at 30 kHz with 16-bit resolution. Action potentials (spikes) were detected by applying a threshold set at −3.5 times the electrode’s root mean square voltage. Single neurons were identified using the k-medoids clustering algorithm, with the optimal number of waveform clusters determined by the gap criterion. (Supplemental Figure 4) Clustering was performed on a space comprised of principal components that collectively accounted for 95% of the waveform variance. Lastly, average FR is calculated as the number of spikes per second for each sorted single neuron. We use the FR instead of raw counts due to randomized trial lengths. Each neuron’s activity is quantified as the average FR across the entire “Go” phase of the trial (2.5-3.5 seconds).

### Empirical Tuning Curves and Surfaces

We represented neuron tuning curves nonparametrically. Let *r*_*t*_ be the firing rate on trial *tt*. Let x∈ {*H, E*} be our two effector Hand and Eye, respectively. Let *C* ∈ {*ME*, S*E*} be the two experimental contexts. For each effector, let 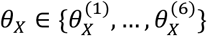 be the angles of the 6 targets and *b*_x_ (*t*) ∈ {1, …,6} be the target index for trial *t*. Let 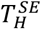, 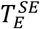, and 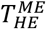 be the set of single effector hand, single effector eye, and multi-effector trials, respectively. We defined single effector tuning curves as

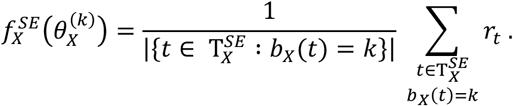

Similarly, we defined the multi-effector tuning surface as

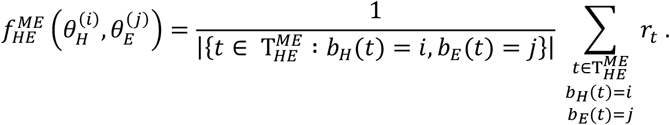

These nonparametric forms allow us to compare the representations with raw data rather than models that approximate the data.

### Additive separability is equivalent to zero cross-partial derivatives

#### Claim

A function is additively separable if and only if it has no cross-partial derivatives

#### Proof

(⇒) Additively separable ⇒ zero cross-partial

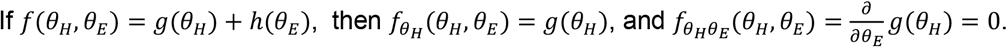

(⇐) Zero cross-partial ⇒ additively separable

Assume 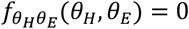. Then for each fixed *θ*_*H*_, the function has derivative

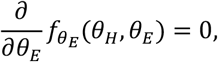

so 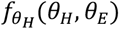 is independent of *θ*_E_ . Hence there exists a function *g*(*θ*_*H*_) such that

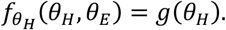

Integrating with respect to *θ*_*H*_ gives us

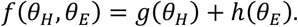

where the “constant of integration” *h* may depend on *θ*_E_. Which is the equation for additive separability.

So,

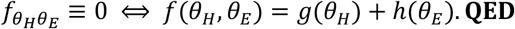

### Separability Calculations

A tuning surface (function) is additively separable if and only if it has no cross-partial derivatives. Cross-partial derivatives are defined as

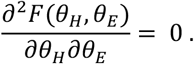

This cross-partial derivative can be calculated numerically on the discrete grid using centered finite differences with circular wrap-around indexing in both angular dimensions. (see Cross Derivative Finite Difference)

Tuning surfaces of neurons which are not mixed selective, are trivially additively separable because they are functions of only one variable. For example,

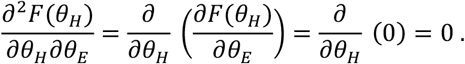

Calculating the cross-derivative of neuron tuning surfaces is useful method when determining the interactions of variables in neural representations. This method benefits from naturally extending to systems with increasing numbers of effectors via the formalism of the Hessian matrix. The Hessian matrix quantifies the local curvature of a function by quantifying its second-order partial derivatives with respect to a pair of variables. A function (*F*) of two variables has a 2x2 Hessian matrix of the form

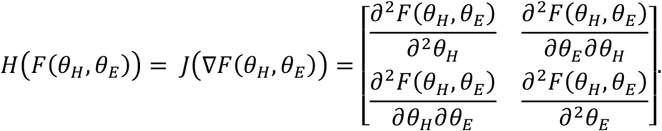

In the context of neuronal representations, each element of a 2×2 Hessian reflects how sharply a neuron’s firing rate changes with respect to hand and eye movements, either independently (diagonal terms) or jointly (off-diagonal terms). If a tuning surface’s Hessian is diagonal over its whole domain, it has no cross-partial derivatives and therefore is additively separable.

Additively separable functions have no cross terms. Therefore, additively separable tuning surfaces can be expressed as a simple sum of independent tuning curves

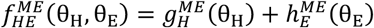

where 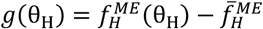 is the centered hand tuning curve averaged over all eye movements and 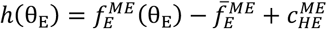 is the centered eye tuning curve averaged over all hand movements plus the ME baseline activity (see Projections and Additive Surfaces).

When calculating all of these cross-derivative quantities, biological noise and measurement error prevent values from being exactly zero. Therefore, cross-derivative values were measured as the root-mean-squared (rms) deviation (*r*_*RMS*_; see Root-Mean-Squared Calculation) of cross-derivative of the tuning surface from zero. These cross-derivative deviations could subsequently be compared to deviations arising from baseline noise.

### Two Pass Smoothing

To reduce high-frequency sampling noise while preserving large-scale curvature structure, smoothing was applied twice sequentially:

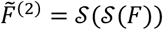

where 𝒮 (⋅) denotes the circular kernel smoothing operation (see Circular Kernel Mean and Variance).

This 2-pass procedure is equivalent to convolution with the kernel twice and results in a smoother effective kernel with reduced variance.

The identical 2-pass smoothing procedure was applied to both empirical and permuted tuning surfaces. Therefore, smoothing does not bias the permutation test (see Separability Permutation Test), as both observed and null statistics are computed under the same estimator.

The rational for this decision is that second derivatives scale like:

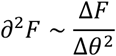

and thus amplify bin-level noise by a factor proportional to 1/Δ*θ*^2^. On a coarse 6-bin grid, this amplification is substantial. The 2-pass smoothing stabilizes derivative estimates while preserving meaningful curvature structure at the scale of behavioral sampling.

### Cross Derivative Finite Difference

When testing additive separability, we quantified the cross-derivative energy of the multi-effector tuning surface 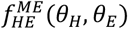. On the discrete 6 × 6 grid, mixed partial derivatives were estimated using centered finite differences with circular wrap-around indexing in both angular dimensions.

Specifically, at grid point (*h, e*),

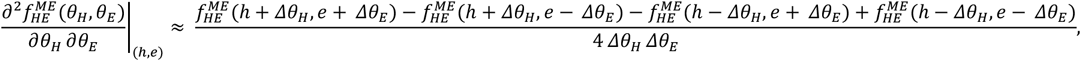

with uniform spacings Δ*θ*_*H*_ and *Δθ*_E_ .

For each neuron we summarized interaction strength by the root-mean-square [Hz/rad^2^] (see Root-Mean- Squared Calculation) across evaluated grid points from a zero reference. Since finite sampling noise ensures a nonzero baseline *r*_RMS_ even under true separability, we constructed a permutation-based null. For each neuron, firing rates were randomly permuted (1000 times) across trials while preserving angular coordinates. The tuning surface, its derivatives, and the RMS statistics were recomputed for each permutation to form a null distribution, and significance was assessed using a right-tailed permutation test. An FDR-BH adjustment was used due to repeated tests for each neuron. Positive test results implied neurons had significant cross-derivatives/variable interactions and negative results implied neurons were additively separable with no variable interactions.

### Root-Mean-Squared Calculation

When comparing the deviation of cross-derivative surface from zero we used root-mean-squared as a metric. For each neuron we evaluated root-mean-square across all grid points,

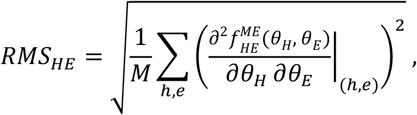

where M is the number of points included.

### Projections and Additive Surfaces

Additive separability enables the transformation of ME tuning surfaces into ME tuning curves as well as the combination of SE tuning curves into SE tuning surfaces. Projections of separable ME tuning surfaces onto individual variables are

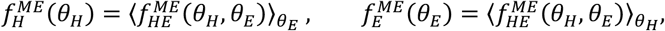

where the symbol ⟨⋅⟩ denotes averaging across the variable’s bins. Centered versions of these projections

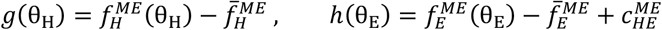

can be understood as tuning curves of each variable in the ME context, where 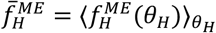 and are 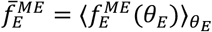 the means (baseline firing rates) of the projections. The constant 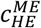 is the grand mean of tuning surface 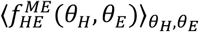. Centering tuning curves prevents double-counting the baseline FRs and the offset 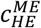 restores the baseline appropriate to the context being modeled.

These ME tuning curves can now be easily compared to their SE tuning curves counter parts using representational similarity metrics (see Representation Similarity Metrics). However, using our equation for additive separability

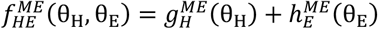

we can reconstruct ME tuning surfaces from our projection based ME tuning curves without loss of information. Such reconstructed ME surfaces take the form

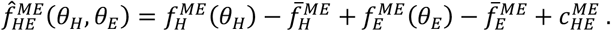

Similar to the reconstructed ME tuning surface, raw empirical SE tuning curves can also be composed into additively compositional tuning surfaces as

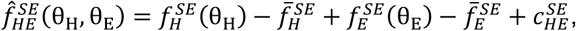

Where 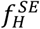 and 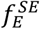 are the SE tuning curves, 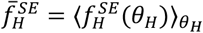 and 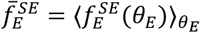 are their means, and 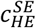 is the predicted mean of the SE tuning surface determined using a weighted average of 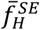 and 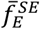 (see Weighted SE Baseline). Centering of the tuning curves 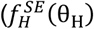 and 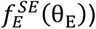 prevents double-counting the baseline FR and the offset 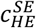 restores the baseline appropriate to the context being modeled. This SE tuning surface allows for comparison of ME tuning surfaces with new tuning surfaces as predicted purely from SE tuning curves.

### Representation Similarity Metrics

Equipped with tuning surfaces and curves in both contexts ({*ME*, S*E*}) we can compare representations with a couple key metrics. Firstly, the preferred direction (PD) for an effector *X* ∈ {*H, E*} and context *C* ∈ {*ME*, S*E*} is the bin of maximal mean response,

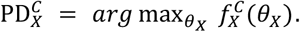

Another important metric is the amplitude of a tuning curve as the difference in firing rate between the peak and trough of the tuning curve:

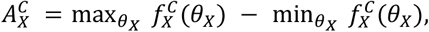

where 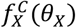 is the tuning curve for each effector *X* and each context *C*. Figure 3A illustrates this definition for the ME data using a light gray segment is centered at 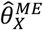 and spans 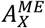. However, the two contexts in *C* have different numbers of trials. The fewer number of points used to calculate the tuning curve in the SE context increases the amplitude variance. To account for such differences, we z-scored the tuning curve amplitudes using a permutation-based amplitude null distribution,

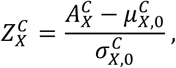

with *µ*_*X*,0_ and *σ*_*X*,0_ being the mean and standard deviation of the amplitude null distribution 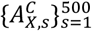. The amplitude null distribution was created by shuffling trial labels (independently for hand and eye) and recomputing the six-bin tuning curve. Each shuffle (*s*) creates permuted amplitude 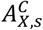 that is used in the null distribution.

### Weighted SE Baseline

Our results demonstrated that ME tuning surfaces had baselines most similar to the baselines of the dominantly encoded effector. This inspired the use of a weighted SE baseline that could predict the baseline of the ME tuning surface from SE tuning curves. This weighted baseline FR was defined as

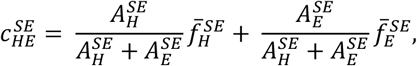

where 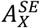 and 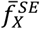 are the neurons’ tuning curves’ amplitudes and means, respectively.

### Proof of Population Separability

In this paper we claim that if single neurons are additively separable then linear combinations (linear projections) of those neurons are also additively separable.

#### Claim

If 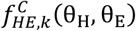 is additively separable ∀*k* ∈ {1, …, *N*} then a linear projection of the population 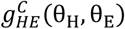 is also additively separable.

#### Proof 1

Let us define a linear combination of neurons as,

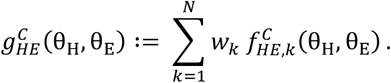

Assuming additive separability of the neurons,

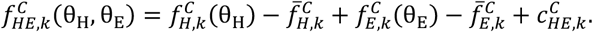

We can then plug this definition of 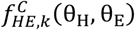 into the linear combination and distribute such that

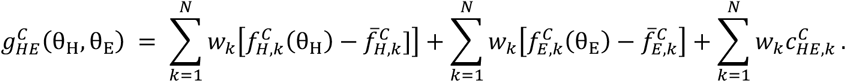

Observing the structure we can simplify each sum on the RHS such that

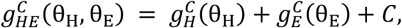

which is additive separability of the linear combination by definition. **QED**

#### Proof 2

Let us define a linear combination of neurons as,

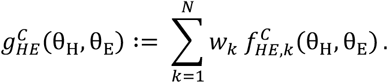

By linearity of differentiation,

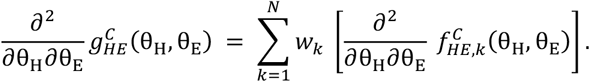

Assuming 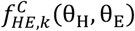 are additively separable, we know their cross derivative is zero. Therefore

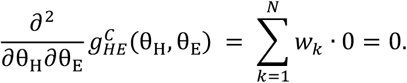

Thus *g* satisfies the necessary and sufficient condition for additive separability. **QED**

### Circular Kernel Mean and Variance

When estimating likelihoods based on nonparametric tuning representations the simplest approach is to compute per-bin means, treating each bin across the curve or grid as independent categories. However, this approach leaves out useful information about the relationship between points. Tuning curves should be continuous, so using information from adjacent bins to smooth, improves bin mean estimates. To do this we standardized the per-neuron FRs and used a kernel-based smoothing estimator. We used a von Mises kernel (*K*) to respect circular geometry, which can be written as

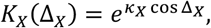

where Δ_*X,t*_ = *d*(*θ*_*X*_, *θ*_*X,t*_ )is a wrapped angular difference and *k*_*X*_ is concentration (bandwidth) controlling smoothness. (small *k* = more smoothing; large *k* = less smoothing) Next, we did 2D or 1D Nadaraya– Watson smoothing [50–52],

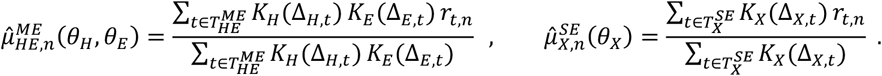

The bandwidth κ was chosen per neuron by 5-fold cross-validation on the training trials (*CCCC*) to minimize the Gaussian negative log-likelihood

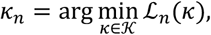

where is 𝒦 = {2,4,8,12, …,32} and

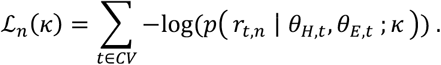

The likelihood *p* (*r*_*ref,n*_ *∣ θ*_*H*_, *θ*_E_ )is defined below (see Bayesian Inference).

When adding the tuning curves together we again did the following:

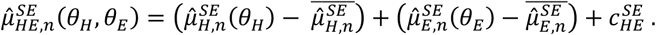

We accompany the mean value calculations with predictive noise models. For the ME surface we calculated global variance as well as local heteroscedastic variance using kernel weights. We define these variance calculation as

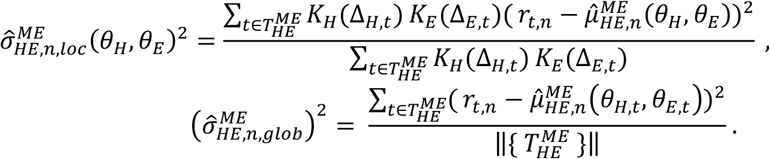

To provide conservative regularization we also perform shrinkage toward each neuron’s global variance

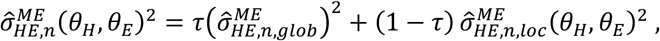

with 0 ≤ *τ* ≤ 1 (we used *ττ* = 0.3). Alternatively, for the SE surfaces we kept the variance calculations independent and simply added them as we did with the means. To assess variance for each independent effector, we calculated

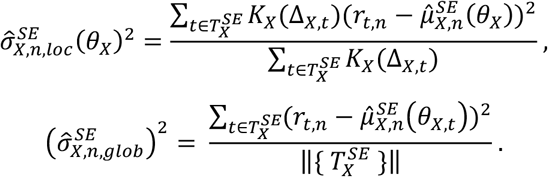

After calculating the variance independently, we also performed shrinkage

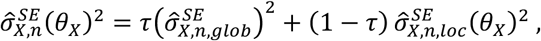

toward each variables global variance, separately (again we used *τ* = 0.3). We then calculated SE surface variance by adding the independent variance values

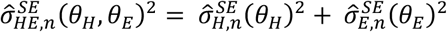

as we did with the mean values. The mean and variance values calculated here are used to specify gaussian likelihoods used for stimulus inference.

### Bayesian Inference

For a single neuron, a tuning surface, given any test trial 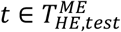 with observed firing vector *r*_*t,n*_, can be turned into a likelihood

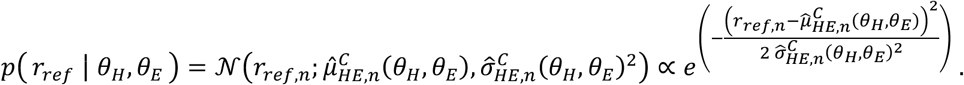

However, we recorded from many neurons that all respond to these movement stimuli. If we assume that neuron responds roughly independently to a given stimulus the population likelihood becomes

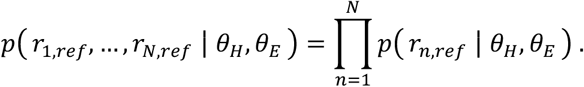

Using Bayes’ equation for inference,

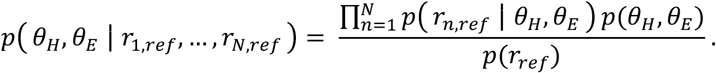

Assuming a uniform prior on movement directions and dropping the marginal distribution for firing rates as a proportionality constant the posterior is proportional to the likelihood,

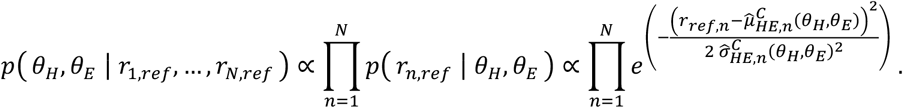

Using this population posterior we performed a maximum a posteriori (MAP) estimate of *θ* _*H*_ & *θ* _*S*_ given observed population firing rates to *r*_1,*ref*_, …, *r*_*N,ref*_. Because of assuming a uniform prior this map estimate was equivalent to a maximum likelihood estimate (MLE). Thus, we equivalently calculated,

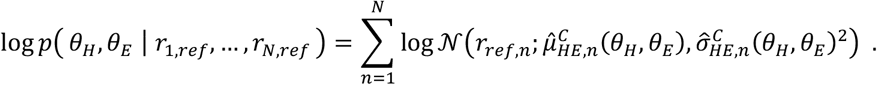

We evaluated *argma*x_*θ*∈*Θ*_ log *p* (*θ*_*H*_, *θ*_E_ *∣ r*_1,*r*ef_, …, *r*_*N,r*ef_ )on all 36 states of *ΘΘ* are returned 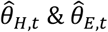.

### Inverted Dropping Curve

To characterize how decoding performance scaled with population size, we constructed inverted dropping curves within each recording session. For each session containing *N* simultaneously recorded neurons, decoding accuracy was evaluated as a function of the number of neurons included in the model.

For a given population size x ∈ {1, …, *N*}, neurons were randomly sampled without replacement from the available pool within that session. For each sampled subset, the decoder was trained and evaluated on the test set. Decoding accuracy was computed as the proportion of correctly classified trials. This procedure was repeated multiple times for each x (with independent random subsets) to reduce sampling variance.

To visualize overall scaling trends across sessions, accuracy values for each sampled subset were pooled across sessions, yielding a set of points {(x, *y*)}. No cross-session normalization of neuron count was applied; instead, curves were evaluated up to the maximum available neuron count in each session.

To summarize the scaling relationship and estimate asymptotic performance, pooled accuracy values were fit with a saturating exponential model of the form

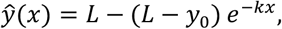

where x is the number of neurons, *y*_0_ is the empirical accuracy at x = 1, *L* > 0 represents the asymptotic performance limit, *k* > 0 controls the rate of saturation.

Parameters *L* and *k* were estimated using least squares. For visualization purposes, exponentials were extrapolated beyond the maximum observed population size using the fitted parameters.

## QUANTIFICATION AND STATISTICAL ANALYSIS

All statistical analyses were performed using custom scripts in MATLAB, and all tests were two-sided unless otherwise noted.

### Weighted Coefficient of Determination

To quantify reconstruction quality across neurons, we compared 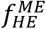 and 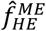 over the common 6×6 grid using a weighted coefficient of determination,

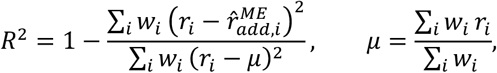

where i indexes hand–eye bin pairs and wi can equal the number of trials contributing to cell i (or wi≡ 1 when coverage is uniform). Weighting guarded against spuriously penalizing sparsely sampled bins.

### Separability Permutation Test

To assess statistical significance of RMS deviation, we constructed a permutation-based null distribution separately for each neuron. Within each neuron firing rates were randomly permuted across trials to create a random ME surface. Subsequently the RMS cross-derivative value was recalculated based on the random ME surface. The random surface, as well as the observed surface, were subject to two pass smoothing to reduce second derivative noise during this test (see Two Pass Smoothing). This process was repeated many times (N=1000 permutations), yielding a neuron-specific null distribution:

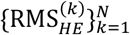

This null preserves the empirical firing-rate distribution and sampling density of each grid bin but destroys any systematic dependence on hand and eye angle.

Then, for each neuron, a one-sided p-value was computed as

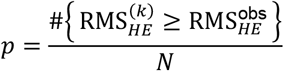

reflecting the probability that a random surface would exhibit interaction curvature at least as large as the observed value.

Because separability was tested across large neuronal populations, we controlled the false discovery rate (FDR) at *q* < 0.05 using the Benjamini–Hochberg procedure. Resulting q-values were used for all statistical inferences.

### Selectivity Permutation Tests and Bootstrapping

Permutation tests were also used to determine the statistical significance of effector-specific tuning (selectivity) based on the representation metrics (see Representation Similarity Metrics). All statistical procedures were fully nonparametric to accommodate heteroscedastic and non-Gaussian firing-rate distributions.

To quantify modulation of a neuron’s firing rate by a given effector, we defined the z-scored tuning amplitude 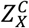, where *X* ∈ {*H, E*} denotes hand or eye and *C* denotes the experimental context (single-effector or multi-effector). Tuning amplitude was computed as the range of the tuning curve across target angles, normalized by the standard deviation of firing rates across trials.

To assess whether observed tuning exceeded chance levels, we employed a nonparametric, permutation-based statistical test. For each neuron and condition, we generated a null distribution 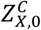 by randomly permuting trial labels across target-angle bins, thereby preserving the marginal firing-rate distribution while destroying any systematic relationship between firing rate and movement direction. A one-sided p-value was then computed as

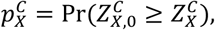

corresponding to the probability that a neuron drawn from the null distribution exhibited tuning amplitude greater than or equal to the observed value.

Because significance testing was performed across large neuronal populations, we controlled the false discovery rate (FDR) at *α* = 0.05 using the Benjamini–Hochberg procedure. Each family of p-values (ME-H, ME-E, SE-H, SE-E) was converted into q-values 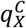, which were used for all selectivity assignments.

During initial analyses, we found that increasing the number of permutations alone did not yield stable q-values across different random seeds, particularly for neurons near threshold. To improve robustness, we therefore embedded the permutation test within a bootstrap procedure. Specifically, we generated *B* = 500 bootstrap resamples by sampling trials with replacement within each condition. For each bootstrap resample *b*, we recomputed the tuning amplitude 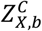, corresponding null distribution, and FDR-corrected q-value 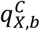.

This procedure produced a distribution of q-values 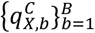 and effect sizes 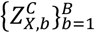. We defined the median q-value

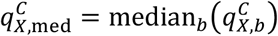

as our final significance metric, and the median tuning amplitude

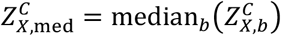

as the corresponding effect size.

Neurons were classified as hand-selective 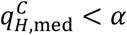, eye-selective if 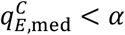, both-selective (mixed-selective) if both were significant, and none-selective otherwise.

## SUPPLEMENTAL INFORMATION

**Document S1. Figures S1–S4**

**Video S1. H-E Coordination Center-out Task, related to Figure 1**

**Video S2. Attempted reaches during the H-E Coordination Center-out Task, related to Figure 1**

## REFERENCES

[1] M. Mishkin, L. G. Ungerleider, and K. A. Macko, Object vision and spatial vision: two cortical pathways, Trends in Neurosciences 6, 414 (1983).

[2] W. P. Medendorp, H. C. Goltz, T. Vilis, and J. D. Crawford, Gaze-Centered Updating of Visual Space in Human Parietal Cortex, J. Neurosci. 23, 6209 (2003).

[3] J. D. Connolly, M. A. Goodale, J. F. DeSouza, R. S. Menon, and T. Vilis, A comparison of frontoparietal fMRI activation during anti-saccades and anti-pointing, J Neurophysiol 84, 1645 (2000).

[4] R. A. Andersen, G. K. Essick, and R. M. Siegel, Encoding of spatial location by posterior parietal neurons, Science 230, 456 (1985).

[5] M. Vesia and J. D. Crawford, Specialization of reach function in human posterior parietal cortex, Exp Brain Res 221, 1 (2012).

[6] A. P. Batista, C. A. Buneo, L. H. Snyder, and R. A. Andersen, Reach Plans in Eye-Centered Coordinates, Science 285, 257 (1999).

[7] C. A. Buneo, M. R. Jarvis, A. P. Batista, and R. A. Andersen, Direct visuomotor transformations for reaching, 416, 5 (2002).

[8] A. Batista, Inner space: Reference frames, Current Biology 12, R380 (2002).

[9] L. H. Snyder, A. P. Batista, and R. A. Andersen, Coding of intention in the posterior parietal cortex, Nature 386, 167 (1997).

[10] L. H. Snyder, K. L. Grieve, P. Brotchie, and R. A. Andersen, Separate body- and world-referenced representations of visual space in parietal cortex, Nature 394, 887 (1998).

[11] R. Q. Quiroga, L. H. Snyder, A. P. Batista, H. Cui, and R. A. Andersen, Movement Intention Is Better Predicted than Attention in the Posterior Parietal Cortex, J. Neurosci. 26, 3615 (2006).

[12] L. H. Snyder, J. L. Calton, A. R. Dickinson, and B. M. Lawrence, Eye-Hand Coordination: Saccades Are Faster When Accompanied by a Coordinated Arm Movement, Journal of Neurophysiology 87, 2279 (2002).

[13] L. H. Snyder, A. P. Batista, and R. A. Andersen, Saccade-Related Activity in the Parietal Reach Region, Journal of Neurophysiology 83, 1099 (2000).

[14] S. W. C. Chang, A. R. Dickinson, and L. H. Snyder, Limb-Specific Representation for Reaching in the Posterior Parietal Cortex, J. Neurosci. 28, 6128 (2008).

[15] S. W. C. Chang, C. Papadimitriou, and L. H. Snyder, Using a Compound Gain Field to Compute a Reach Plan, Neuron 64, 744 (2009).

[16] E. Salinas and T. J. Sejnowski, Gain Modulation in the Central Nervous System: Where Behavior, Neurophysiology, and Computation Meet, (2010).

[17] M. Brozovic, L. F. Abbott, and R. A. Andersen, Mechanism of gain modulation at single neuron and network levels, J Comput Neurosci 25, 158 (2008).

[18] L. R. Bremner and R. A. Andersen, Temporal Analysis of Reference Frames in Parietal Cortex Area 5d during Reach Planning, J. Neurosci. 34, 5273 (2014).

[19] L. R. Bremner and R. A. Andersen, Coding of the Reach Vector in Parietal Area 5d, Neuron 75, 342 (2012).

[20] B. Pesaran, M. J. Nelson, and R. A. Andersen, Dorsal Premotor Neurons Encode the Relative Position of the Hand, Eye, and Goal during Reach Planning, Neuron 51, 125 (2006).

[21] E. J. Hwang, M. Hauschild, M. Wilke, and R. A. Andersen, Spatial and Temporal Eye-Hand Coordination Relies on the Parietal Reach Region, Journal of Neuroscience 34, 12884 (2014).

[22] J. D. Crawford, W. P. Medendorp, and J. J. Marotta, Spatial Transformations for Eye–Hand Coordination, Journal of Neurophysiology 92, 10 (2004).

[23] P. N. Sabes, B. Breznen, and R. A. Andersen, Parietal Representation of Object-Based Saccades, Journal of Neurophysiology 88, 1815 (2002).

[24] A. Pouget and T. J. Sejnowski, Spatial transformations in the parietal cortex using basis functions, J. Cognitive Neuroscience 9, 222 (1997).

[25] A. Pouget and L. H. Snyder, Computational approaches to sensorimotor transformations, Nat Neurosci 3, 1192 (2000).

[26] R. A. Andersen and C. A. Buneo, Intentional maps in posterior parietal cortex, Annu Rev Neurosci 25, 189 (2002).

[27] Y. E. Cohen and R. A. Andersen, A common reference frame for movement plans in the posterior parietal cortex, Nat Rev Neurosci 3, 553 (2002).

[28] M. C. Bushnell, M. E. Goldberg, and D. L. Robinson, Behavioral enhancement of visual responses in monkey cerebral cortex. I. Modulation in posterior parietal cortex related to selective visual attention, J Neurophysiol 46, 755 (1981).

[29] V. N. Christopoulos, J. Bonaiuto, I. Kagan, and R. A. Andersen, Inactivation of Parietal Reach Region Affects Reaching But Not Saccade Choices in Internally Guided Decisions, J Neurosci 35, 11719 (2015).

[30] J. C. Culham and K. F. Valyear, Human parietal cortex in action, Current Opinion in Neurobiology 16, 205 (2006).

[31] R. A. Andersen, K. N. Andersen, E. Hwang, and M. Hauschild, Optic ataxia: from Balint’s syndrome to the parietal reach region, Neuron 81, 967 (2014).

[32] A. Z. Khan, L. Pisella, Y. Rossetti, A. Vighetto, and J. D. Crawford, Impairment of Gaze-centered Updating of Reach Targets in Bilateral Parietal–Occipital Damaged Patients, Cereb Cortex 15, 1547 (2005).

[33] M.-M. Mesulam, Spatial attention and neglect: parietal, frontal and cingulate contributions to the mental representation and attentional targeting of salient extrapersonal events, Philos Trans R Soc Lond B Biol Sci 354, 1325 (1999).

[34] T. Aflalo et al., Decoding motor imagery from the posterior parietal cortex of a tetraplegic human, Science 348, 906 (2015).

[35] K. K. Noneman and J. Patrick Mayo, Decoding Continuous Tracking Eye Movements from Cortical Spiking Activity, Int. J. Neur. Syst. 35, 2450070 (2025).

[36] M. F. Land, Eye movements and the control of actions in everyday life, Progress in Retinal and Eye Research 25, 296 (2006).

[37] O. Donchin, A. Gribova, O. Steinberg, H. Bergman, and E. Vaadia, Primary motor cortex is involved in bimanual coordination, Nature 395, 274 (1998).

[38] P. J. Ifft, S. Shokur, Z. Li, M. A. Lebedev, and M. A. L. Nicolelis, A Brain-Machine Interface Enables Bimanual Arm Movements in Monkeys, Sci. Transl. Med. 5, (2013).

[39] J. E. Downey, K. M. Quick, N. Schwed, J. M. Weiss, G. F. Wittenberg, M. L. Boninger, and J. L. Collinger, The Motor Cortex Has Independent Representations for Ipsilateral and Contralateral Arm Movements But Correlated Representations for Grasping, Cerebral Cortex 30, 5400 (2020).

[40] F. R. Willett, D. R. Deo, D. T. Avansino, P. Rezaii, L. R. Hochberg, J. M. Henderson, and K. V. Shenoy, Hand Knob Area of Premotor Cortex Represents the Whole Body in a Compositional Way, Cell 181, 396 (2020).

[41] N. P. Shah et al., Pseudo-linear Summation explains Neural Geometry of Multi-finger Movements in Human Premotor Cortex, bioRxiv 2023.10.11.561982 (2023).

[42] D. R. Deo, F. R. Willett, D. T. Avansino, L. R. Hochberg, J. M. Henderson, and K. V. Shenoy, Brain control of bimanual movement enabled by recurrent neural networks, Sci Rep 14, 1598 (2024).

[43] M. Abeles, M. Diesmann, T. Flash, T. Geisel, M. Herrmann, and M. Teicher, Compositionality in neural control: an interdisciplinary study of scribbling movements in primates, Front. Comput. Neurosci. 7, (2013).

[44] M. S. Willsey, N. P. Shah, D. T. Avansino, N. V. Hahn, R. M. Jamiolkowski, F. B. Kamdar, L. R. Hochberg, F. R. Willett, and J. M. Henderson, A Real-Time, High-Performance Brain-Computer Interface for Finger Decoding and Quadcopter Control.

[45] X. Zou et al., Control of a Commercially Available Vehicle by a Tetraplegic Human Using a Brain-Computer Interface, https://arxiv.org/abs/2508.11805v2.

[46] C. Guan, T. Aflalo, K. Kadlec, J. Gámez De Leon, E. R. Rosario, A. Bari, N. Pouratian, and R. A. Andersen, Decoding and geometry of ten finger movements in human posterior parietal cortex and motor cortex, J. Neural Eng. 20, 036020 (2023).

[47] C. Y. Zhang, T. Aflalo, B. Revechkis, E. R. Rosario, D. Ouellette, N. Pouratian, and R. A. Andersen, Partially Mixed Selectivity in Human Posterior Parietal Association Cortex, Neuron 95, 3 (2017).

[48] K. M. Tye, E. K. Miller, F. H. Taschbach, M. K. Benna, M. Rigotti, and S. Fusi, Mixed selectivity: Cellular computations for complexity, Neuron 112, 2289 (2024).

[49] K. Kadlec, J. Gamez, T. Aflalo, C. Guan, E. R. Rosario, K. Pejsa, A. Bari, N. Pouratian, and R. Andersen, Distinct Patterns of Whole-Body Coding in Human Motor Cortex and Posterior Parietal Cortex.

[50] E. A. Nadaraya, On Estimating Regression, Theory Probab. Appl. 9, 141 (1964).

[51] G. S. Watson, Smooth Regression Analysis, Sankhy?: The Indian Journal of Statistics, Series A (1961-2002) 26, 359 (1964).

[52] K. V. Mardia and P. E. Jupp, Directional Statistics (J. Wiley, Chichester ; New York, 2000).

