## Supplemental Document 1 for "The Compositional Encoding of Hand-Eye Coordinated Movements for Single Neurons in the Posterior Parietal Cortex"

### Supplemental Information

Nikos Mynhier,[1,5,*] Jorge Gamez,[1] Kelsie Pejsa,[1]
Ausaf Bari,[3,4] Richard M. Murray,[1,2,4] and Richard Andersen[1,4,**]
[1]Division of Biology and Biological Engineering, California Institute of Technology, Pasadena, CA, USA
[2]Division of Engineering and Applied Science, California Institute of Technology, Pasadena, CA, USA
[3]Department of Neurosurgery, David Geffen School of Medicine at UCLA, Los Angeles, CA, USA
[4]Senior author
[5]Lead contact

#### FIGURE S1

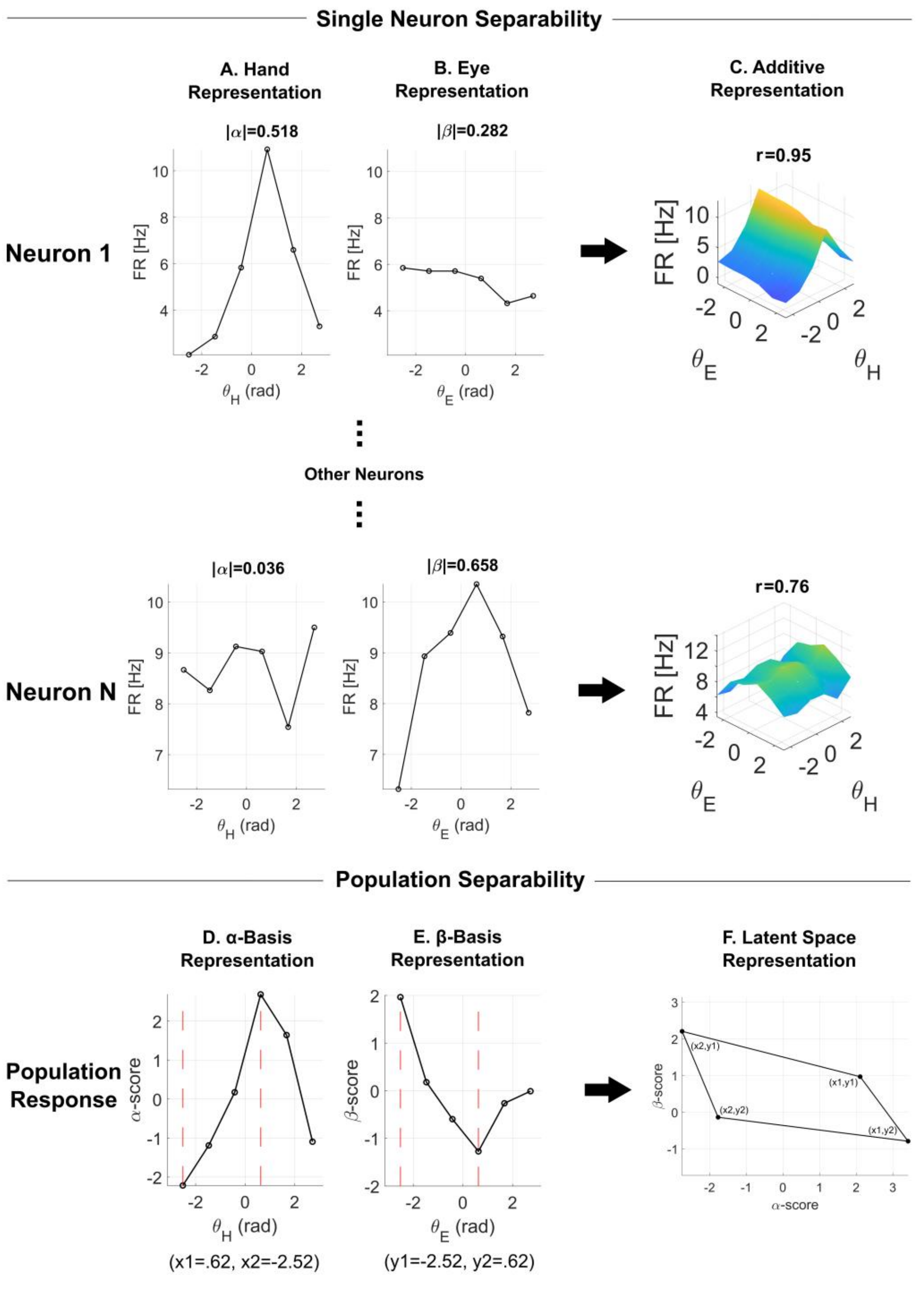


**Figure S1. Single-neuron and population additivity tested with a demixed additive basis (DAB).**
**All data presented here is taken from a single representative clinical session. (A–C) Single neuron activity. For each neuron we compute mean firing rates on a (6x6) hand–eye grid and test additive separability. (A) Marginal ME hand tunings of single neurons; (B) Marginal ME eye tunings of single neurons. The neurons' absolute DAB loadings ($|\alpha_n|$) and ($|\beta_n|$) are reported above the tuning curves. (C) Additive ME surface ($\hat{f}_{HE}^{ME}(\theta_H, \theta_E)$); the correlation of the additive surface (corr($f_{HE}^{ME}, \hat{f}_{HE}^{ME}$)) to the raw ME surfaces is shown above each additive surface.**
**(D–F) Projected population activity. We standardize each neuron across conditions (per-neuron z-score), row-center the resulting matrix, and construct a demixed additive basis $\{\alpha,\beta\} \subset R^N$ by SVD of the population marginals. (See Methods) (D) α-basis score vs hand direction (averaged across eye directions). (E) β-basis score vs eye direction. Vertical red lines mark the peak and trough of panel D (x1,x2) and panel E (y1,y2). (F) Four projected points $\{s(x_i,y_j), t(x_i,y_j)\}$ for $((x_i, y_j) \in \{x1, x2\} \times \{y1, y2\})$ in the population latent space that form a parallelogram.**

**FIGURE S2**

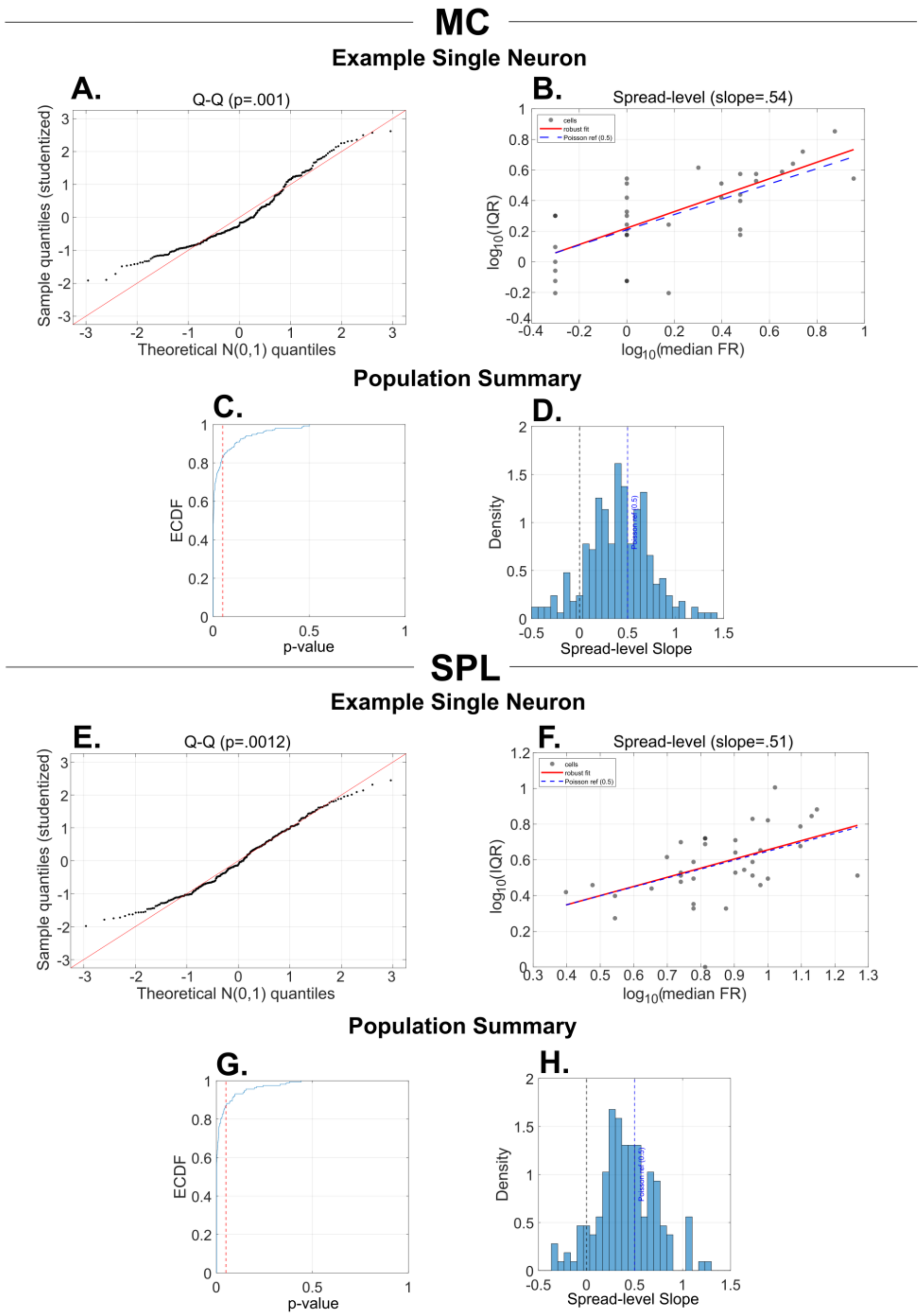


**Figure S2. Non-normality and heteroskedasticity of single neurons in MC and SPL**

**Panels A–D show results for the MC and Panels E–H show results for the PPC. For each brain area the top two plots represent single-neuron results and bottom two plots represent population level results. (A) Q–Q plots of studentized residuals of an example neuron against a N(0,1) reference (red line). Residuals are defined trial-wise as $z_{ijt} = (r_{ijt} - \bar{r}_{ij})/s_{ij}$ , where $\bar{r}_{ij}$ and $s_{ij}$ are the mean and SD within each hand–eye combination $(i, j)$. Clear curvature/tail excess indicates non-Gaussian errors even after within-cell scaling. The reported $p$-value is from a Lilliefors test on the pooled studentized residuals. (B) Spread–level plot for an example neuron. $log_{10}(IQR_{ij})$ versus $log_{10}(median_{ij})$ for each hand-eye combination $(i, j)$. The solid line is a least squares linear fit for the scattered data. The blue dashed line represents mean-variance ratio ($\sigma \propto \mu^{1/2}$) of a Poisson process (slope=.5). Positive slopes in general indicate heteroskedasticity. (C) ECDF of Lilliefors p-values across all neurons in the MC. The vertical dashed line marks p=0.05. Neurons with p-values less than .05 are significantly different from a normal distribution. (D) Histograms of spread–level slopes for all neurons in the MC. The dotted vertical lines mark 0 (no mean–variance relation) and 0.5 (Poisson mean-variance relation). Distributions centered above zero indicate heteroskedasticity. (E-H) Same plots as A-D but for neurons in the SPL.**

#### FIGURE S3

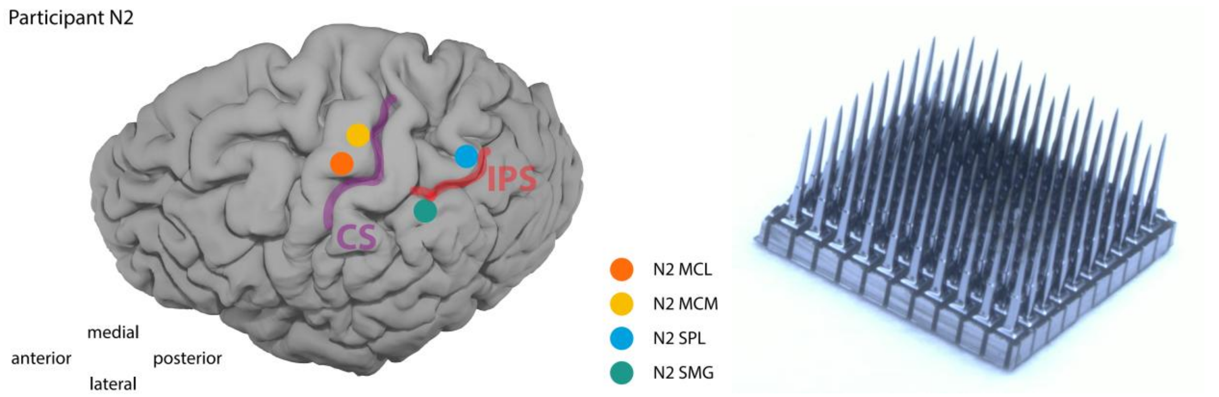


**Figure S3: Implant Locations and Neuroport Array.**
**Anatomical locations of participant RD's implants. Zoomed in photo of the Neuroport Utah electrode array.**

#### FIGURE S4

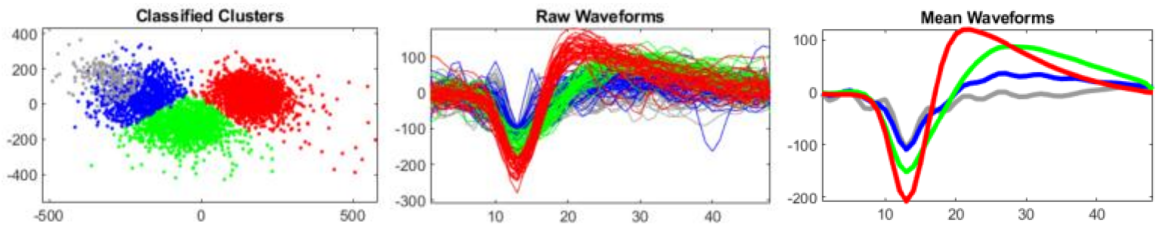


**Figure S4: Spike Sorting**
**Standard output of our spike sorting algorithm. First plot is a scatter plot of all recorded spikes from a single electrode projected into the first two principal components of waveform space. This example shows three clusters in waveform space corresponding to three unique single units. The second plot shows all recorded spike waveforms color coded by their cluster. The third plot shows the mean waveform of each cluster.**
